# Timely sleep coupling: spindle-slow wave synchrony is linked to early amyloid-β burden and predicts memory decline

**DOI:** 10.1101/2022.03.15.484463

**Authors:** Daphne Chylinski, Maxime Van Egroo, Justinas Narbutas, Vincenzo Muto, Mohamed A. Bahri, Christian Berthomier, Eric Salmon, Christine Bastin, Christophe Phillips, Fabienne Collette, Pierre Maquet, Julie Carrier, Jean Marc Lina, Gilles Vandewalle

## Abstract

Sleep alteration is a hallmark of ageing and emerges as a risk factor for Alzheimer’s disease (AD). While the fine-tuned coalescence of sleep microstructure elements may influence age-related cognitive trajectories, its association with AD processes is not fully established. Here, we investigated whether the coupling of spindles and slow waves is associated with early amyloid-beta (Aβ) brain burden, a hallmark of AD neuropathology, and cognitive change over 2 years in 100 healthy individuals in late-midlife (50-70y; 68 women). We found that, in contrast to other sleep metrics, earlier occurrence of spindles on slow-depolarisation slow waves is associated with higher medial prefrontal cortex Aβ burden (p=0.014, r^2^_β*_=0.06), and is predictive of greater longitudinal memory decline (p=0.032, r^2^_β*_=0.07). These findings unravel early links between sleep, AD-related processes and cognition and suggest that altered coupling of sleep microstructure elements, key to its mnesic function, contributes to poorer brain and cognitive trajectories in ageing.

## INTRODUCTION

Alterations in sleep quality are typical of the ageing process with a more fragmented and less intense (or shallower) sleep detected as early as the fifth decade of life^1^. Beyond healthy ageing, alterations in sleep are predictive of the risk of developing Alzheimer’s disease (AD) over the next 5 to 10 years^2,3^. Similarly, sleep disorders such as insomnia and obstructive sleep apnoea syndrome are associated with increased odds for AD diagnosis^4,5^. Brain burdens of aggregated amyloid-β (Aβ) and tau proteins, hallmarks of AD pathophysiology, have been linked with a reduced sleep intensity, as indexed by the overall production of slow waves during sleep, but also to worse objective sleep efficiency and subjective sleep quality, in healthy and cognitively normal older individuals aged > 70y^6–11^. Sleep alteration may in turn contribute to the aggregation of AD proteins: experimental sleep deprivation and sleep fragmentation (disturbance in the production of slow wave during sleep) lead to increased concentration of Aβ in the cerebrospinal fluid (CSF), both in animal models and in healthy human populations^7,9,12,13^. Overall, a bidirectional detrimental relationship between sleep quality and the neuropathology of AD is emerging in the literature. Sleep may therefore constitute a modifiable risk factor which one could act upon to prevent or delay the neuropathological processes associated to AD and favour successful cognitive trajectories over the lifespan^14–16^. Hitherto, however, sleep is not yet widely recognised as an independent risk factor for AD and the mechanistic associations between sleep and early AD neuropathology are not yet fully established.

Sleep microstructure elements, such as sleep spindles and slow waves (SW), are essential correlates of cognitive function of sleep, as higher densities of both elements during post-learning sleep have been linked to a better overnight memory consolidation^17–21^. Furthermore, SW activity (i.e. a power measure combining the density and amplitude of sleep SW over sleep cycles in the 0.75 to 4 Hz band) was reported to modulate the regression between prefrontal cortex Aβ burden and a lower overnight memory consolidation in cognitively normal older individuals^6^. The fine-tuned coupling of spindles and SW has further been reported to be altered in ageing, with an earlier occurrence of the spindle relative to the SW depolarisation phase in the older compared to younger individuals, and to predict overnight memory retention^22^. Whether this spindle-SW coupling in ageing is associated to AD-pathological processes and cognitive trajectories is not fully established, however. A study in 31 individuals, aged around 75y, found a link between the brain deposit of tau protein and the coupling of spindles and SW^23^. By contrast, another research failed to find a link between this coupling and Aβ brain burden^24^. Here, we argue that, on top of potential statistical power issues, the difficulty to detect this link may be due to the fact that the assessments were carried out late over the lifespan (i.e. > 70y), when subtle associations may be masked by concurrent brain alterations, and by the heterogeneity of sleep SW.

The low frequency oscillations of the electroencephalography (EEG) have been divided in slow oscillations (≤ 1 Hz) and delta waves (1-4 Hz) for decades in humans, notably based on the theoretical framework of the generation of SW^25–27^. As one of the main features of the SW resides in their transition from down-to up-state, reflecting synchronised depolarisation, a recent work proposed the transition frequency of the down-to-up state as a way of distinguishing between *slow* and *fast* switcher SWs^28^. Compared to young adults, on top of exhibiting a typical overall lower density of SW, older individuals reportedly exhibit higher probabilities of producing slow as compared to fast switcher SWs, providing important insights into age-related changes in sleep microstructure.

Investigating the coupling of spindles and SW, appropriately split between the slow and fast switchers, in late middle-aged healthy adults may be the best approach to gain insight into the biology underlying the early relationship between sleep and AD-related processes. In a longitudinal study, we therefore tested whether the coupling of spindles with the slow and the fast switcher SWs is associated with the early brain burden of Aβ and memory performance in a large sample (N=100) of healthy and cognitively normal individuals in late midlife (50-70y). We recorded habitual sleep in these individuals devoid of sleep disorders of both sexes (59.5±5y; 68 woman) under EEG and extracted the density and coupling of spindle and fast and slow switcher SWs over frontal derivations. The burden of Aβ was assessed using Positron Emission Tomography (PET) tracers (^[18F]^Flutemetamol, N=96; ^[18F]^Florbetapir, N=4) over the medial prefrontal areas, known to be an early site for Aβ deposits and the most important generator of SWs during sleep^6,29–31^. Performance to the Mnemonic Similarity Task (MST), a memory task highly sensitive to early signs of cognitive decline^32,33^, was assessed in all participants concomitantly to EEG and PET measurements (N=100) as well as at follow-up, 2 years later, in a substantial subsample (N=66). We hypothesised that our large sample of individuals, positioned relatively early in the ageing process, would allow to detect subtle associations between the impaired fine-tuned coupling of spindles and slow and fast switcher SWs, and both the early Aβ burden and memory performance decline over 2 years.

## RESULTS

### Slow but not fast switcher SW show preferential coupling with spindles

We first assessed whether spindles were specifically associated with a particular phase on the down-to-up state transition of the SWs, considering the two types of SWs (slow and fast switchers). Of the 341.836 detected slow switcher SWs, 75.910 co-occurred with a spindle (22%); while of the 78.235 fast switcher SWs, 26.912 (34%) were found to co-occur with a spindle. Regarding spindles, 563.928 spindles were detected over all the recordings, of which 102.822 were coupled to a SW (18%), 75.910 to slow switcher SWs (13%) and 26.912 to fast switcher SWs (5%). Statistical analysis with Watson’s U^2^ test showed that the distribution of spindle onset was significantly different between slow and fast switcher SWs (**Figure 1B**) (U^2^ = 71.143, p <0.001). Indeed, spindles were preferentially anchored only onto slow switchers, as spindles were most often initiated on the ascending phase of the depolarised state of the slow switcher SWs, while no such preferred coupling was detected for fast switcher SWs (**Figure 1C**). As the slow switcher SWs are the only SW to exhibit a preferential coupling with spindles, they remained our main focus of interest for the remaining analyses.

**Figure 1.**
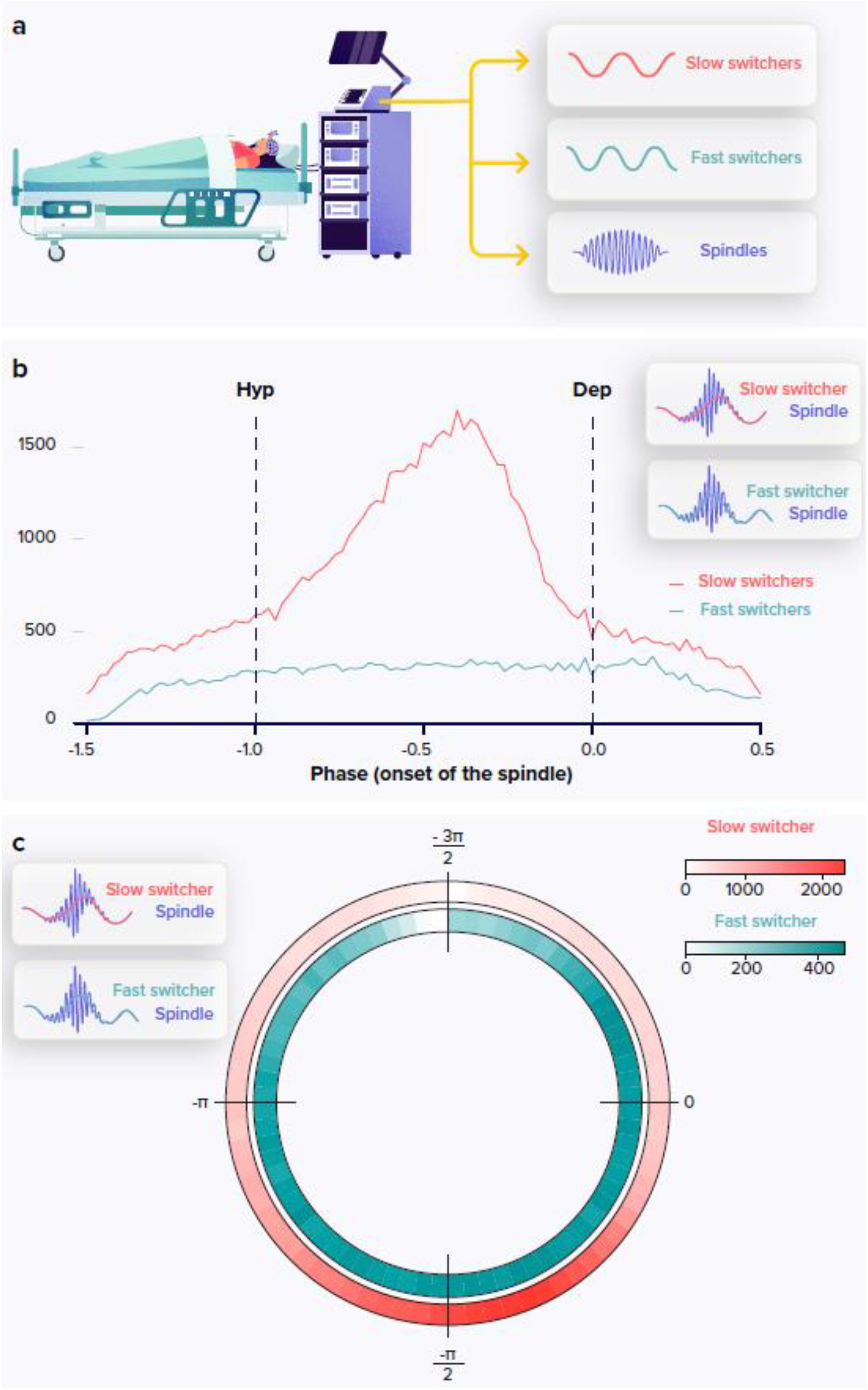
Following a screening night and a regular sleep-wake schedule for 1 week, the participants (N=100; age +-; 68 women) slept in the lab at their habitual times under EEG recording. We extracted the density and coupling of spindles and fast and slow switcher SWs over frontal derivations during N2 and N3 sleep stage from EEG recordings (panel A). Analysis of the anchoring of the spindles onto the SWs showed a preferential coupling phase only for slow (red) but not fast switcher SWs (light blue) (the y axis represents the number of spindles starting at a specific SW phase) (panel B). Circular representation of the anchoring of the spindles onto the SW phase: −π/2 represents the hyperpolarisation (down state), 0 represents the depolarisation of the SW. Heatmap represents the density of the spindles with their onset on specific slow wave phase in 5° bins across all participant nights, for slow (red) and fast switcher SWs (light blue)(panel C).

### Spindle onset on slow switcher SW is linked to prefrontal Aβ burden

Statistical analysis revealed that the anchoring of the spindle onset onto slow switcher SWs was significantly linked to the burden of Aβ over the medial prefrontal cortex (MPFC) (**Figure 2A**) (main effect of Aβ PET uptake: F_1,96_=6.2, **p=0.014, r**^**2**^_**β\***_**=0.06**), where earlier onset of the spindle relative to the SW phase was associated with higher Aβ PET uptake (**Figure 2B**). This effect was detected while controlling for the differences between sexes (main effect of sex: F_1,96_=5.01, p=0.028, r^2^_β*_=0.05), as a later spindle onset was found in men relative to women, and controlling for age (main effect of age, F_1,96_=0.04, p=0.85). We assessed the specificity of this association and show that the anchoring of the spindle onset onto fast switcher SWs was not linked to the Aβ burden over the MPFC (main effect of Aβ PET uptake: F_1,96_=0.20, p=0.65), after controlling for age (main effect of age: F_1,96_=0.41, p=0.53) and sex (main effect of sex: F_1,96_=0.48, p=0.49) (**Figure 2C**). In addition, the density of slow switcher SWs (main effect of Aβ PET uptake: F_1,96_=0.14, p=0.71) or of spindles (main effect of Aβ PET uptake: F_1,96_=0.09, p=0.76) was not associated with the MPFC Aβ burden (**Figure 2D-E**), further reinforcing the idea that it is the coupling of sleep microstructure elements that matters rather than their individual occurrence. Likewise, unlike previous reports^6,23^, we did not find any association between the MPFC Aβ burden and several characteristics of the SWs (SW density - per min of NREM sleep - F_1,95_=1.18, p=0.28; spindle density – per min of NREM sleep - F_1,95_=0.06, p= 0.80; cumulated power generated in the delta band - or slow wave energy (SWE) - F_1,95_=0.84, p=0.36; the proportion of slower oscillations (0.5-1 Hz) over the delta power (1.25-4 Hz) - SO/delta proportion - F_1,95_=2.82, p=0.10), after correcting for age, sex and total sleep time (*SW density*: age: F_1,95_=5.49, p=0.02, r^2^_β*_=0.05; sex: F_1,95_=10.26, p=0.002, r^2^_β*_= 0.1; TST: F_1,95_=0.5, p=0.48; *spindle density*: age: F_1,95_=0.56, p=0.46, sex: F_1,95_=0.42, p=0.52, TST: F_1,95_=0.36, p=0.55; *SWE*: age: F_1,95_=4.2, p=0.04, r^2^_β*_=0.04; sex: F_1,95_=6.63, p=0.01; TST: F_1,95_=0.9, p=0.34; *SO/delta proportion*: age: F_1,95_=,3.28 p=0.07, sex: F_1,95_=0.0, p=0.96, TST: F_1,95_=2.7, p=0.10) (**Suppl. Fig. 1**).

**Figure 2.**
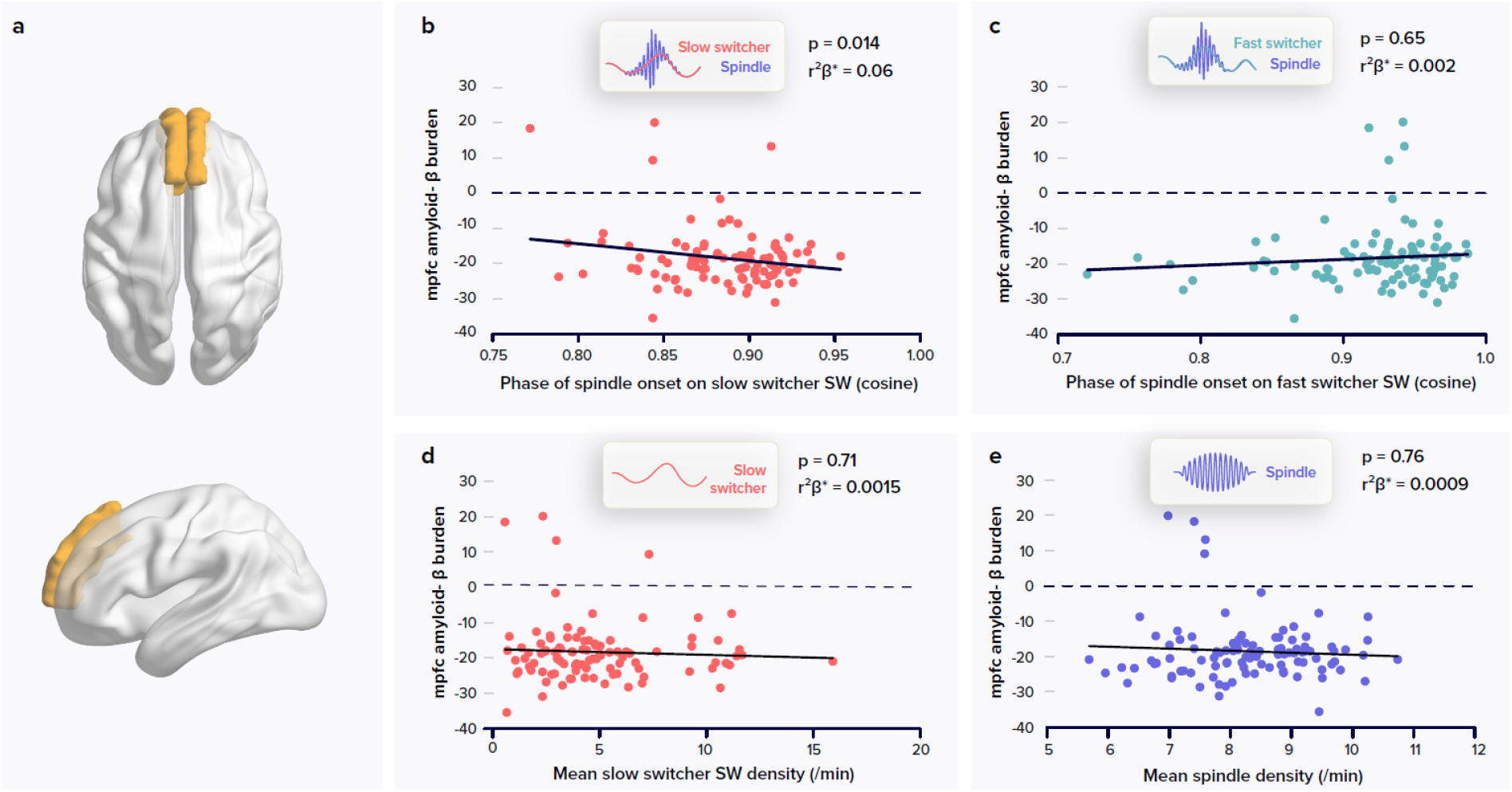
Relationships between spindles and slow waves metrics and the amyloid-β (Aβ) burden. PET signal uptake was measured over the medial prefrontal cortex (MPFC) depicted in yellow (panel A). Significant negative association between the MPFC Aβ burden and spindle-slow switcher SW coupling (panel B). No association between the MPFC Aβ burden and spindle-fast switcher SW coupling (panel C). No association between the MPFC Aβ burden and slow switcher SW density (panel D). No association between the MPFC Aβ burden and spindle density (panel E). P values and r^2^β* were computed from the GLMMs referred to in the text. Simple regressions were used only for a visual display and do not substitute the GLMM outputs. We used the cosine value of the phase of coupling in the GLMMs (see methods).

To test the apparent difference in the association between the coupling of spindles onto slow and fast switcher SWs with the accumulation of Aβ protein, we further computed a statistical model with spindle-SW coupling as the dependent variable, while including the SW type together with the Aβ MPFC burden as independent variables. Statistical analysis yielded a significant interaction between the burden of Aβ in the MPFC and the type of SWs (Aβ burden by SW type interaction: F_1,96_=7.05, **p=0.009; r**^**2**^_**β\***_**=0.07)**, while *post-hoc* tests indicated that the link between the coupling of the spindle onto the SW and the MPFC Aβ burden was significant for the slow switcher type (t_149.8_=-2.00, **p=0.047**) and not for the fast switcher type (t_149.8_=0.56, p=0.57). This finding reinforces the idea that slow and fast switcher SWs constitute distinct realisations of NREM oscillations that are differently associated with brain aggregation of Aβ during the ageing process. This has likely contributed to previous failures to detect links between the coupling of spindles and SW and the deposit of Aβ. In fact, when testing the association between the coupling of spindles and SWs, irrespective of the type of SWs, and PET Aβ burden over the MPFC, the statistical analysis only yields a weak negative association between the phase of the coupling of the spindle onto the SW and Aβ burden (main effect of Aβ uptake: F_1,96_=3.96, **p=0.049, r**^**2**^_**β\***_**=0.04**; main effect of sex: F_1,96_=4.33, p=0.04, r^2^_β*_=0.04; main effect of age: F_1,96_=0.05, p=0.83), which could arguably go undetected in a smaller or different sample.

### Slow switchers spindle phase coupling is associated to memory change over two years

We tested whether the coupling of spindles with slow switcher SWs was associated with memory performance and its decline over 2 years using the mnemonic similarity task (MST) (**Figure 3A**). The MST consists in a pattern separation task targetting the ability to distinguish between highly resembling memory events, a hippocampus dependent task which is very sensitive to early cognitive decline^32,33^. Across the sample, we observed an overall decline in performance between the baseline and follow-up performance at the MST (t_65_ = 2.19, **p=0.032**). We found no significant link between the coupling of the spindles onto both SW types and the performance on the task at baseline (i.e. assessed at the same time as the sleep measures) (slow switchers: *main effect of spindle-SW coupling*: F_1,96_=0.26, p=0.61; main effect of age: F_1,96_=0.36, p=0.55, main effect of sex: F_1,96_=0.41, p=0.52, main effect of education: F_1,96_=0.47, p=0.50; fast switchers: *main effect of spindle-SW coupling*: F_1,96_=0.02, p=0.88; main effect of age: F_1,96_=0.34, p=0.56, main effect of sex: F_1,96_=0.54, p=0.46, main effect of education: F_1,96_=0.40, p=0.53) (**Figure 3B-C**). By contrast, statistical analyses revealed a significant negative link between the relative change in memory performance and the phase of spindle anchoring onto slow switcher SWs, indicating that an earlier spindle onset is predictive of a memory worsening over two years (main effect of spindle-slow switcher SW coupling: F_1,61_=4.80, **p=0.032, r**^**2**^_**β\***_**=0.07**), after correcting for age (main effect of age: F_1,61_=0.25, p=0.62), sex (main effect of sex: F_1,61_=0.20, p=0.66) and education (main effect of education: F_1,61_=0.25, p=0.62) (**Figure 3D-E**). No such association was detected when considering spindle coupling to fast switcher SWs (main effect of spindle-fast switcher SW coupling: F_1,61_=2.51, p=0.12; main effect of age: F_1,61_=0.33, p=0.57, main effect of sex: F_1,61_=0.18, p=0.68, main effect of education: F_1,61_=1.11, p=0.30). Further statistical analyses show that the memory performance change over the 2-year is not significantly related to the MPFC Aβ burden (main effect of Aβ burden: F_1,60_=2.33, p=0.13; main effect of age: F_1,60_=1.27, p=0.26, main effect of sex: F_1,60_=0.03, p=0.87, main effect of education: F_1,60_=0.41, p=0.53).

**Figure 3.**
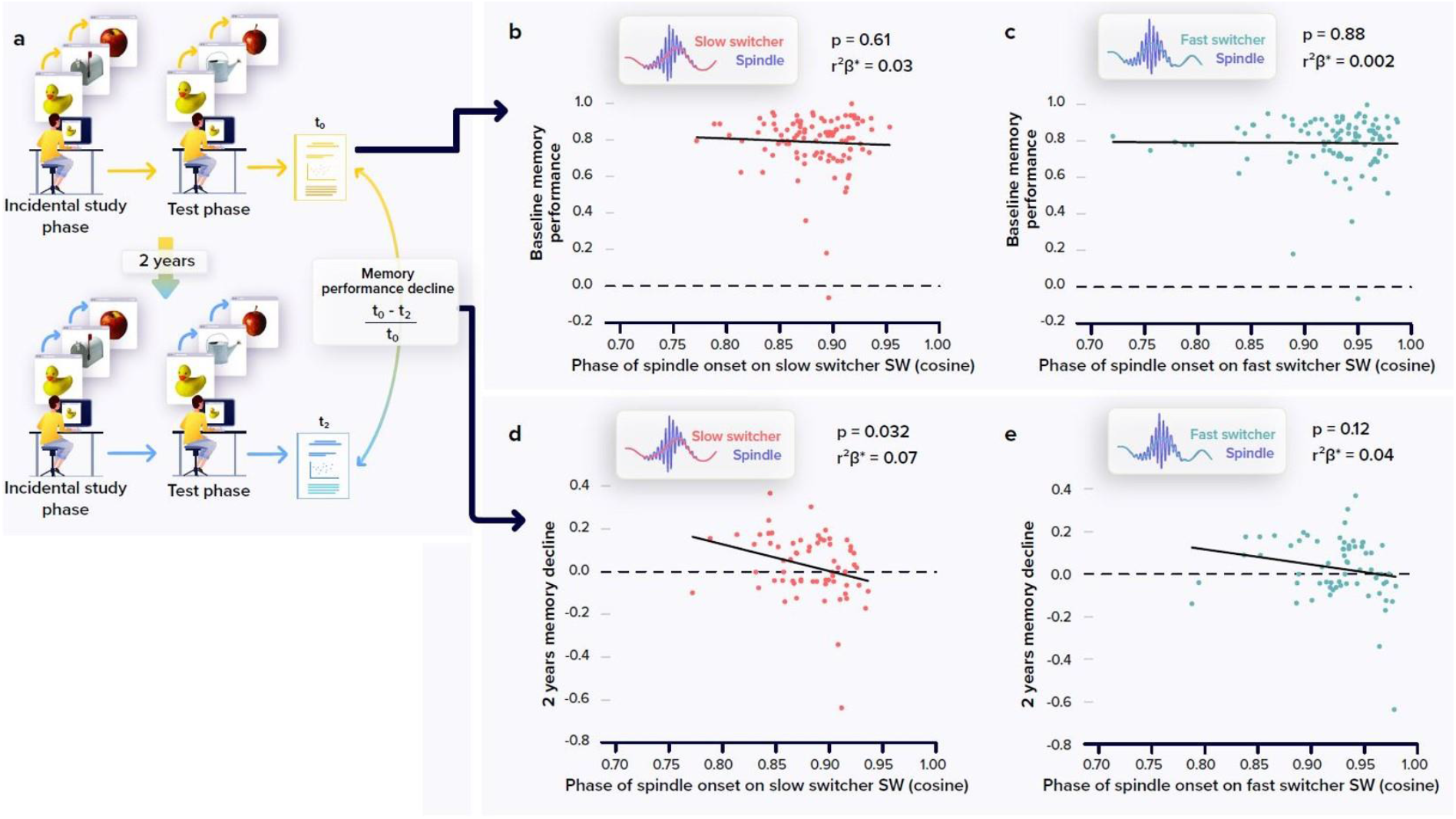
Relationships between memory performance and coupling between spindles and SWs. Memory performance was assessed through the Mnemonic Similarity Task (MST) where participants have to recognized previously encoded images in series of new or lure images (see methods) (panel A). No association between the baseline MST performance and spindle-slow switcher SW coupling (panel B). No association between the baseline MST performance and spindle-fast switcher SW coupling (panel C). Significant negative association between the 2 years relative change in MST performance and spindle-slow switcher SW coupling (panel D). No association between the 2 years relative changes in MST performance and spindle-fast switcher SW coupling (panel D). P values and r^2^β* were computed from GLMMs referred to in the text. Simple regressions were used only for a visual display and do not substitute the GLMM outputs. We used the cosine value of the phase of coupling in the GLMMs (see methods).

**Figure 4.**
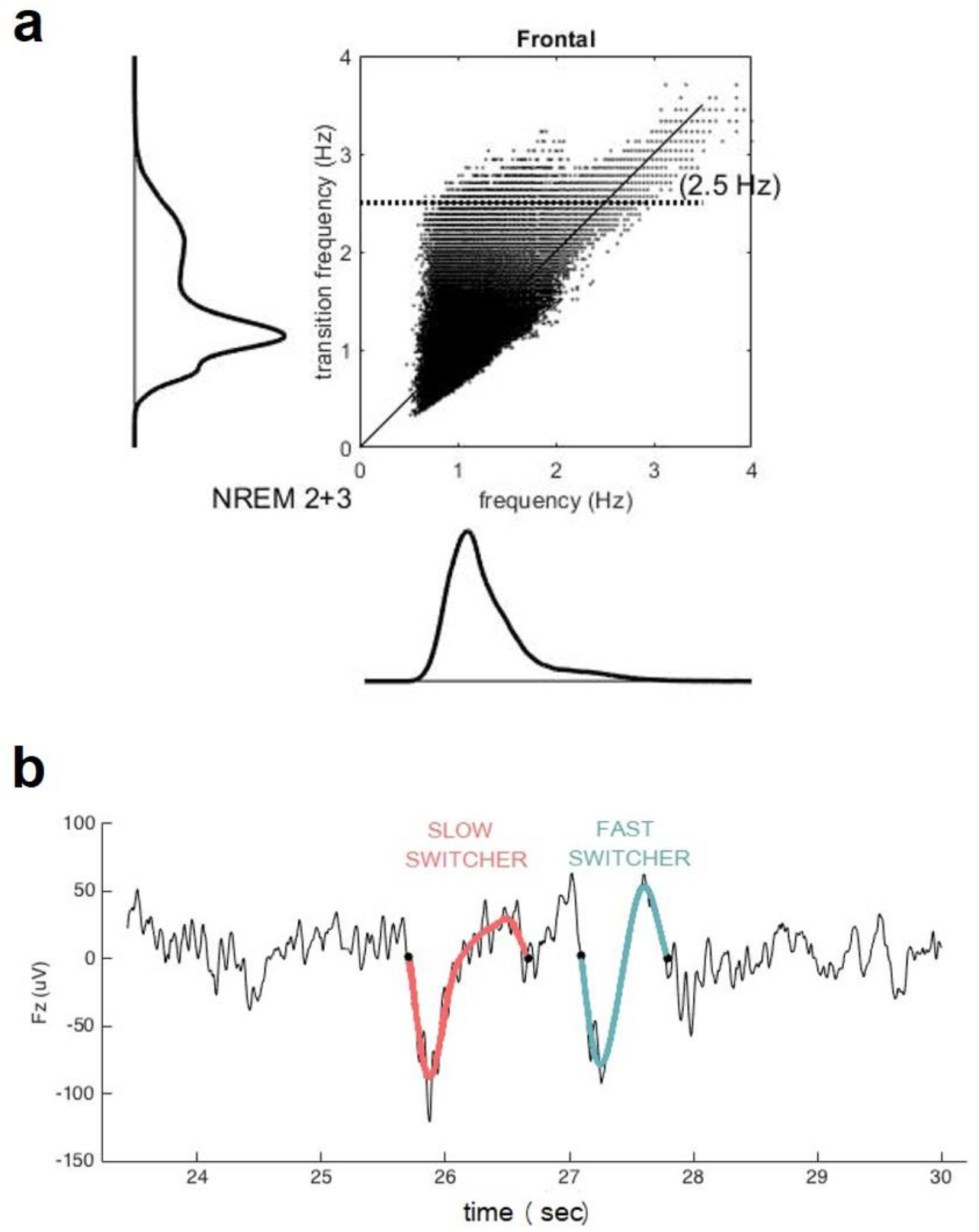
Distribution of the mean frequency of the slow waves (x axis) versus their transition frequency (y axis) for both NREM2 and NREM3 sleep stages (panel A) in the entire study sample. One ccan observe a double distribution of the frequency of transition but not in the overall frequency. Example of slow switcher (red) and fast switcher SW (light blue) extracted from the EEG signal.

## DISCUSSION

In order to unravel early associations between the microstructure of sleep and the burden of Aβ in the brain, and their cognitive implications, we collected polysomnography, PET and behavioral data in a relatively large sample of individuals without cognitive impairments or sleep disorders. To this end, we recruited individuals in late middle age (50-70y), that could in most cases only present limited age-related alterations in sleep and accumulation of Aβ protein in the brain^34^. We investigated whether the coupling of spindles onto SWs, showing a slower and a faster frequency of transition from the down to the up states (slow and fast switcher SWs) was associated to the accumulation of Aβ over the medial prefrontal cortex. We further probed whether the coupling of spindles onto SWs was associated with the performance to a sensitive memory test, assessed at the time of the sleep and PET recordings and, longitudinally, 2 years later. The coupling of spindles onto the slow, but not the fast, switcher SWs was signficantly associated with the Aβ PET signal assessed over the MPFC. Moreover, this coupling between spindles and slow switcher SW was significantly linked to the memory performance change detected 2 years after the initial assessement. Overall, our results provide compelling evidence that the link between sleep and the accumulation of Aβ over the MPFC, an early AD-related brain features, involves the precise and timely coupling of two key elements of NREM sleep, spindles and SWs, and that this coupling bears a predictive value for the subsequent decline in memory performance. Our study does not indicate, at least not in these healthy and relatively young older adults, that the amount of spindles or SW generated overnight is associated with the accumulation of Aβ over the MPFC.

Sleep SWs provide a readout of the homeostatic sleep pressure and are more prevalent at the beginning relative to the end of the night^35^. In addition, both the density of spindles and SWs have separately been related to overnight consolidation of memory^21,36^. They actively take part in information transfer from hippocampic to neocortical networks and in synaptic plasticity^17,21^. Recent research has put forward the importance of their precise phase coupling during sleep, and reported an age-related difference in that coupling^22^. In the younger individuals, spindles tend to reach their maximum around the cortical up-state of the SWs, whereas in older individuals, spindles occur earlier on the depolarisation phase of the SWs. This earlier coupling between spindles and SWs is related to a poorer overnight memory retention^22^, suggesting a sub-optimal neuronal interplay for the exchange of information during sleep in older individuals. In line with a presumed suboptimal coupling between spindles and SWs, we find that, when controlling for age, individuals for which the spindles occur earlier during the transition phase of the slow switcher SWs, show higher Aβ burden over the MPFC and a worse memory change over time. We did not find any significant effect of the age of participants on the phase of the coupling between spindles and SWs. This is likely due in part to the restricted age range of our participants, but also to the variability existing between the individuals in the changes they undergo in their sleep during ageing. Importantly, the relationship we observed between the phase of the spindle onset onto the SW and the MPFC Aβ burden shows the same directionality (i.e. earlier spindles onto SW) as the changes previously reported in older individuals.

SWs were previously characterised according to the frequency of their transition from the down-to the up-state, which reflects the relative synchronisation of the depolarisation of the neurons when generating a SW^28^. Beyond the well-known decrease in the production of SWs in aging, the slow switcher SWs were relatively preserved in the older individuals compared to the fast switcher SWs. This finding suggests that the two populations of SWs constitute distinct elements of sleep microstructure. Three present results confirm that the two types of SW –slow and fast switchers – behave differentially. First, sleep spindles show a preferential coupling only with the transition period from down-to-up state of the slow switcher SWs, while spindles occuring concomittantly to fast switchers SWs do not occur at a specific phase of the SWs. Furthermore, only the coupling of spindles onto slow switcher SWs was significantly associated to the early accumulation of Aβ in the brain. Finally, only the coupling of spindles and slow switcher SWs was predictive of the memory change after 2 years. Our results support that slow switcher SWs, the type previously reported to be relatively spared during aging^28^, is important to the development of AD-related pathological changes, at least in the form of Aβ protein accumulation, and for the subsequent development of subtle alteration in the cognitive abilities, leastways over the memory domain.

Sleep spindles are considered as thalamic events. They are generated through the interplay between the inhibitory reticular nucleus and the excitatory thalamocortical neurons, which project to the cortical neurons that feedback in turns to the thalamus^17^. In contrast, SWs are intrinsically cortical. They consist in the spontaneous alternations between down (hyperpolarized) and up (depolarized) neuronal states^37^. The SWs undergo, however, an influence from the thalamus reticular nucleus that contributes to the synchonization of distant neuronal populations^26,37^. Hence, although the exact neurophysiological origin of the functional associations between spindles and SWs remain to be established, the thalamus reticular neurons could arguably be involved. Our results would therefore indicate that the early aggregation of Aβ protein in the medial prefrontal cortex could disturb the thalamocortical interplay driving the coalescence of spindles and SWs. This chronic disturbance would then trigger the cognitive changes detectable after 2 years. The cross-sectional nature of our imaging data does not, however, allow to make inferences regarding the directionality of the relationship between the spindle-slow switcher SW coupling and the burden of Aβ protein. Evidence accumulates to show that if AD-neuropathological hallmarks can affect the quality of sleep, sleep can also impinge onto these hallmarks^14,38^. Specifically relevant in the context of this study, the occurence of the SWs has been associated with a transient increase in the glymphatic flow related to local variations in neuromodulator concentrations^39^. It is therefore possible that the changes in the density of SWs occurring during ageing affect both their coupling with spindles and the early accumulation of Aβ protein.

We found that the density of either spindles or SWs is not related to the early accumulation of the Aβ protein in the medial prefrontal cortex. This suggests a specific role for the coupling of both elements of sleep microstructure. Moreover and as previously reported in an intermediate analysis of a subsample of the same study^40^, we do not find significant associations between more macroscopic measures, which are typically used to characterize sleep, and the accumulation of Aβ protein. The cumulated power of the oscillations generated in the delta band (i.e. the slow wave energy - SWE), and the proportion of slower oscillations over the delta power were not associated with PET measures. This finding argues against the idea that these rougher measures of sleep disruption constitute the earliest manifestations of the association between sleep and AD-related processes and contrasts with previous reports in a smaller sample of individuals older (> 70y) than our sample. Altogether, these discrepancies reinforce the idea that the alteration in the microstructure of sleep, consisting in the coupling of the spindles onto a specific subpopulation of SWs, as reported here, but also in the occurrence of micro-arousals during sleep we previously reported based on the same sample^41^, shows a prior association with AD-related processes compared with the amount of slow brain oscillations generated during overnight sleep. The latter may only be significantly associated at a later age, when the pathophysiological changes are already more substantial. In addition, the coupling of spindles onto slow switcher SWs is predictive of the future change in memory performance. Sleep microstructure could therefore constitute a promising early marker of future cognitive and brain ageing trajectory^41^. We did not evaluate whether distinct links between the SW types and slow and fast spindles are observed. As some reports describe that fast spindles are rather coupled to the up-state of the SWs and slow spindles tend to occur on the waning depolarisation phase of the SWs^42^, we could hypothesise that the associations we observe are probably rather driven by fast spindles. Future investigations are, however, needed to confirm this hypothesis

One should bear in mind the potential limitations of our study. First, although we collected data in a relatively large sample, we may have insufficient power to detect some associations with other sleep measures. One can nevertheless frame our findings in relative terms such that association between spindle-SWs coupling and the early accumulation of Aβ protein is at least stronger than the association with the coupling of spindles onto fast switchers SW, the density of SWs and spindles, the SWE, etc. Also, the longitudinal aspect of our study is relatively short-termed and only concerned the performance to a sensitive mnesic task while it did not include sleep EEG and the PET assessments. Further studies should evaluate the predictive value of such parameters on longer longitudinal protocols, and the evolution of the sleep EEG and the PET parameters as well as their generalisability over other precociously impacted cognitive abilities. Finally, given that our protocol does not include manipulation of the coupling of the spindles onto the SWs, it precludes any inference on the causality of one aspect onto the other.

Together, our findings reveal that the timely occurrence of spindles onto a specific type of SWs showing a relative preservation in ageing seems to play a determining role in ageing trajectory, both at the cognitive level and with regards to structural brain integrity. These findings may help to unravel early links between sleep, AD-related pathophysiology and cognitive trajectories in ageing and warrants future clinical trials attempting at manipulating sleep microstructure or Aβ protein accumulation.

## MATERIALS AND METHODS

### Study design and participants

101 healthy participants aged from 50 to 70 y (68 women; mean ± SD = 59.4 ± 5.3 y) were enrolled between 15 June 2016 and 2 October 2019 for a multi-modal cross-sectional study taking place at the GIGA-Cyclotron Research Centre/In Vivo Imaging of the University of Liège (Cognitive fitness in aging – COFITAGE – study) which has already led to several scientific publications [e.g. ^41,43^]. One participant was excluded from analyses due to lack of PET imaging data. The exclusion criteria were as follows: clinical symptoms of cognitive impairment (Mattis Dementia Rating Scale >130 ; Mini-Mental State Evaluation (MMSE) >27) ; recent history psychiatric history or severe brain trauma ; self-reported or clinically diagnosed sleep disorder ; Body Mass Index (BMI) ≤18 and ≥29; use of medication affecting sleep or the central nervous system; smoking; excessive alcohol (>14 units/week) or caffeine (>5 cups/day) consumption; shift work in the 6 months or transmeridian travel in the 2 months preceding the study. All participants gave their written informed consent prior to their participation. The study was registered with EudraCT 2016-001436-35. All procedures were approved by the Hospital-Faculty Ethics Committee of ULiège. All participants signed an informed consent prior to participating in the study.

**Table 1:** Sample characteristics

|  | Baseline (N=100) | Follow up (N=66) |
| --- | --- | --- |
| Sex | 68 ♀ / 32 ♂ | 44 ♀ / 22 ♂ |
| Age (years) | 59.4 ± 5.3 [50-69] | 59.9 ± 5.4 [50-69] |
| Education (years) | 15.2 ± 3.0 [9 - 25] | 14.9 ± 3.3 [9-25] |
| Total sleep time (TST) (minutes, EEG) | 392.8 ± 45.9 [229 – 495.5] | 390.4 ± 45.9 [264.0 – 495.5] |
| Time spent in N1 sleep stage (% of TST, EEG) | 6.2 ± 2.8 [0.6 - 15.6] | 6.4 ± 3.0 [0.6 – 15.6] |
| Time spent in N2 sleep stage (% of TST, EEG) | 51.6 ± 8.9 [31.4 - 75.7] | 50.3 ± 8.5 [32.8 – 75.7] |
| Time spent in N3 sleep stage (% of TST, EEG) | 19.2 ± 6.4 [7.2 - 38.3] | 19.7 ± 6.5 [8.2 – 38.3] |
| Time spent in REM sleep (% of TST, EEG) | 23.1 ± 6.8 [6.5 – 39.8] | 23.6 ± 7.4 [6.5 – 39.8] |
| Mean SW density (number/minute of N2/N3) | 7.1 ± 4.3 [0.8 – 19.2] | 6.9 ± 4.0 [1.0 – 19.2] |
| Slow switchers | 4.9 ± 3.1 [0.6 - 15.9] | 4.9 ± 3.1 [0.6 – 15.9] |
| Fast switchers | $2.1 \pm 1.6$ [0.1 - 8.8] | $2.0 \pm 1.3$ [0.3 – 5.7] |
| Mean SW amplitude (μV) | $101.5 \pm 12.4$ [76.0 - 128.2] | $101.0 \pm 12.3$ [76.0 – 125.2] |
| Slow switchers | $104.5 \pm 13.7$ [77.7 - 131.3] | $104.0 \pm 13.5$ [77.7 – 130.5] |
| Fast switchers | $94.0 \pm 11.0$ [73.0 - 130.1] | $93.0 \pm 10.1$ [73.5 – 113.7] |
| Mean SW frequency | $1.2 \pm 0.1$ [1.0 - 1.6] | $1.2 \pm 0.1$ [1.1 – 1.6] |
| Slow switchers | $1.1 \pm 0.1$ [1.0 - 1.3] | $1.1 \pm 0.1$ [1.0 – 1.2] |
| Fast switchers | $1.5 \pm 0.1$ [1.3 - 1.8] | $1.5 \pm 0.1$ [1.3 – 1.8] |
| Mean SW transition frequency | $1.4 \pm 0.1$ [1.1 - 1.8] | $1.4 \pm 0.1$ [1.1 – 1.8] |
| Slow switchers | $1.1 \pm 0.0$ [1.0 - 1.2] | $1.1 \pm 0.0$ [1.0 – 1.2] |
| Fast switchers | $2.0 \pm 0.1$ [1.9 - 2.2] | $2.0 \pm 0.1$ [1.9 – 2.2] |
| Mean spindle density (number/minute of N2/N3) | $8.3 \pm 1.1$ [5.7 – 10.7] | $8.1 \pm 1.0$ [6.0 – 10.2] |

*BMI, alcohol consumption, anxiety & depression scores as well as sleepiness levels of the sample can be found* *in* <sup>41</sup>

### Sleep assessment

A first night of sleep was recorded at the laboratory under full polysomnography to avoid potential first night effects and exclude volunteers with sleep apnoea (AHI ≥15/h). A second night of sleep was recorded with electroencephalography (EEG), following one week of regular sleep-wake schedule based on each participant’s preferred bed and wake up time (compliance was verified by actimetry and sleep diary - Actiwatch©, Cambridge Neurotechnology, UK). Sleep was recorded with N7000 amplifiers (EMBLA, Natus, Planegg, Germany). The recording comprised 11 EEG derivations, placed according to the 10-20 system (F3, Fz, F4; C3, Cz, C4; P3, Pz, P4; O1, O2), 2 bipolar electrooculogram (EOGs), and 2 bipolar submental electromyogram (EMG) electrodes. Sampling was set at 200 Hz, and the signal was re-referenced to the mean of the two mastoids. Recordings were scored for sleep stages in 30s windows using a validated automatic algorithm (ASEEGA, Physip, Paris, France) ^44,45^. Automatic arousal and artefact detection ^46,47^ was performed in order to remove EEG segments containing artefacts and arousals from further analysis.

### Slow waves and spindle detection

Only the frontal electrodes were considered because the frontal cortex is an early site showing Aβ deposit and is the primary generator of the SWs during sleep^6,29–31^ as well as to facilitate interpretations of future large-scale studies using headband EEG restricted to frontal electrodes^8^. SWs were automatically detected during N2 and N3 epochs of NREM sleep devoid of artefacts/arousals >5s long, using a previously developed algorithm ^48^. Data were first band-filtered between 0.3 and 4.0Hz with a linear phase Finite Impulse Response (FIR) filter. Following recent work, SW detection criteria were adapted for age and sex ^48^: peak to peak amplitude ≥70μV (resp. ≥60.5μV) and negative amplitude ≤ −37μV (resp. ≤ −32μV) was used for women (resp. for men), instead of the standard ≥75μV and ≤ - 40μV). The duration of the negative deflection had to fit in the range 125-1500ms, and the duration of the positive deflection could not exceed 1000ms. The SWs were sorted according to their transition frequency^28^ (inverse of the duration between the hyperpolarised and depolarised state) into either slow or fast switchers (the critical value for distinguishing between the two types being the intersection between two gaussians, around 1.2Hz^28^). Sleep spindles were also automatically detected over the same N2 and N3 epochs with a previously published method ^49–51^. The EEG signal was bandpass filtered between 10 and 16Hz with a linear phase finite impulse response filter (−3dB at 10 and 16Hz). After detection of SW and spindles, analysis of their coincidence was performed. A coincidence was defined as a co-occurrence of both the ignition and the maximum amplitude of a spindle over the phase of a slow wave. This criterion was used on slow and fast switchers.

### MRI data

Quantitative multi-parametric MRI acquisition was performed on a 3-Tesla MR scanner (Siemens MAGNETOM Prisma, Siemens Healthineers, Erlangen, Germany). Quantitative maps were obtained by combining the images using different parameters sensitive to distinct tissue properties. The multi-parameter mapping was based on multi-echo 3D fast low angle shot at 1 mm isotropic resolution^52^. This included three datasets with T1, proton density (PD), and magnetization transfer (MT)–weighted contrasts imposed by the choice of the flip angle (FA = 6° for PD & MT, 21° for T1) and the application of an additional off-resonance Gaussian-shaped RF pulse for the MT-weighted acquisition. MRI multi-parameter maps were processed with the hMRI toolbox^53^ (http://hmri.info) and SPM12 (Welcome Trust Centre for Neuroimaging, London, UK) to obtain notably a quantitative MT, which was segmented into grey matter, white matter, and CSF using unified segmentation^54^. Flow-field deformation parameters obtained from DARTEL spatial normalisation of the individual MT maps were applied to the averaged co-registered PET images^55^. The volumes of interest were determined using the automated anatomical labelling (AAL) atlas^56^.

### PET-scan

Aβ PET imaging was performed using [^18^F]Flutemetamol, except for 3 volunteers for which [^18^F]Florbetapir was used. PET-scans were performed on an ECAT EXACT+ HR scanner (Siemens, Erlangen, Germany). Participants received a single dose of the radioligand in the antecubital vein (target dose 185±10% MBq); image acquisition started 85min after the injection and consisted of 4 frames of 5 minutes, followed by a 10 minutes transmission scan using ^68^Ge line sources. Images were reconstructed using a filtered back-projection algorithm including corrections for the measured attenuation, dead time, random events, and scatter using standard software (Siemens ECAT – HR + V7.1, Siemens/CTI Knoxville, TN, USA). Individual PET average images were produced using all frames and were then manually reoriented according to MT-weighted structural MRI volumes and coregistered to the individual space structural MT map. Standardised uptake value ratio (SUVR) was computed using the whole cerebellum as reference region^57^. As images were acquired using 2 different radioligands, their SUVR values were converted into Centiloid Units^57^ (the validation of the procedure in our sample was previously published^58^). The Aβ burden was averaged over a mask covering the medial prefrontal cortex previously reported to undergo the earliest aggregation sites for Aβ pathology^34^.

### Cognitive assessments

As part of an extensive neuropsychological assessment, participants were administered the Mnemonic Similarity Task (MST) ^59^, a visual recognition memory task. After an incidental encoding phase during which participants were randomly presented 128 common objects for a period of 2s, and were instructed to determine whether the object presented on the screen was rather an ‘indoor’ or ‘outdoor’ item, the recognition memory phase consisted in the presentation of 192 objects (64 old, presented previously – target items; 64 similar but not identical to the previously presented stimuli – lure; and 64 new objects – foil items). In this phase, participants were instructed to determine whether the presented object was new (foil), previously presented (old), or similar but not perfectly identical (lure). For statistical analyses, the recognition memory score was used (RM), computed as the difference between the rate of calling a target item “old” minus the rate of calling a foil item “old” [P(“old”|target)- P(“old”|foil)] ^32,43^.

The MST was administered at two timepoints: the first time, the day preceding the baseline night, during a cognitive evaluation performed ∼ 6.5h before habitual bedtime. The second neuropsychological evaluation was carried out ∼24 months after the first one (mean 767±54 days). The memory decline score was computed as the baseline performance minus the follow-up performance, divided by the baseline performance, so that a higher score indicates a higher decline over the 2 years.

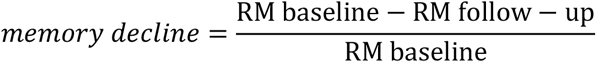

### Statistics

Statistical analyses were performed using Generalised Linear Mixed Models (GLMMs) in SAS 9.4 (SAS Institute, Cary, NC). The distribution of dependent variables was verified in MATLAB 2013a and the GLMMs were adapted accordingly. Subject was treated as a random factor and each model was corrected for age and sex effects. Kenward-Roger’s correction was used to determine the degrees of freedom. Cook’s distance was used to assess the potential presence of outliers driving the associations, and as values ranged below 0.45 no datapoint was excluded from the analyses (a Cook’s distant > 1 is typically considered to reflect outlier value). Our main analysis concerned the coupling between SW types and spindles, and as two analyses were performed (one with slow switcher and one with fast switcher SW), the significance threshold is set at p<0.025 for these analysis, to account for multiple comparisons. The reader should note that we performed a separate statistical test with Aβ burden as dependent variable with both SW-spindle coupling and SW type as independent variables, which confirmed the associations reported in the separate models. The remaining analysis were exploratory as they arise from the main analyses and do not require correction for multiple comparisons. Semi-partial r^2^ (r^2^_β*_) values were computed to estimate the effect sizes of significant fixed effects and statistical trends in all GLMMs^60^. P-values in post-hoc contrasts (difference of least square means) were adjusted for multiple testing using Tukey’s procedure. Watson’s non-parametric two-sample U^2^ test for circular-normal data was performed in MATLAB 2019 to assess the difference between the distribution of spindle onset on the phase of slow waves for slow and fast switcher SW. For analyses using the phase of spindle onset on the slow waves, as the phase of all subjects were in the quarter between the zero crossing (−π/2) and depolarisation, the cosine of the phase was used instead of the phase, in order to perform linear statistics. Statistics with the phase yielded the same results.

Optimal sensitivity and power analyses in GLMM remains under investigation [e.g. ^61^]. We nevertheless computed a prior sensitivity analysis to get an indication of the minimum detectable effect size in our main analyses given our sample size. According to G*Power 3 (version 3.1.9.4)^62^ taking into account a power of .8, an error rate α of .025 (corrected for 2 tests), a sample size of 100 allowed us to detect small effect sizes r > .29 (2-sided; absolute values; confidence interval: .1 – .46; R^2^ > .08, R^2^ confidence interval: .01 – .21) within a linear multiple regression framework including 1 tested predictor (Aβ) and 2 covariates (age, sex).

## Acknowledgements

We thank M. Blanpain, P. Cardone, M. Cerasuolo, E. Lambot, P. Ghaemmaghami, C. Hagelstein, S. Laloux, E. Balteau, A. Claes, C. Degueldre, B. Herbillon, P. Hawotte, B. Lauricella, A. Lesoine, A. Luxen, X. Pepin, E. Tezel, D. Marzoli, C. Schmidt and P. Villar González for their help in different steps of the study. This work was supported by Fonds National de la Recherche Scientifique (FRS-FNRS, FRSM 3.4516.11, Belgium), Actions de Recherche Concertées (ARC SLEEPDEM 17/27-09) of the Fédération Wallonie-Bruxelles, University of Liège (ULiège), Fondation Simone et Pierre Clerdent, European Regional Development Fund (ERDF, Radiomed Project) and the Canadian Institutes of Health Research (CIHR) (grant number 190750). [18F]Flutemetamol doses were provided and cost covered by GE Healthcare Ltd (Little Chalfont, UK) as part of an investigator sponsored study (ISS290) agreement. This agreement had no influence on the protocol and results of the study reported here. M.V.E., C.Bastin, F.C., C.P., and G.V. are/were supported by the F.R.S.-FNRS Belgium.

## Authors contributions

Study concept and design: E.S., P.M., C.P., C.Bastin, F.C. and G.V. Data acquisition: D.C., M.V.E. J.N., V.M., C.Bastin, F.C., G.V. Data analysis and interpretation: all authors. D.C. and G.V. drafted the first version of the manuscript. All authors revised the manuscript and had final responsibility for the decision to submit for publication.

## References

1. Carrier, J. et al. Sleep slow wave changes during the middle years of life. Eur. J. Neurosci. 33, 758–766 (2011).

2. Lim, A. S. P., Kowgier, M., Yu, L., Buchman, A. S. & Bennett, D. A. Sleep Fragmentation and the Risk of Incident Alzheimer’s Disease and Cognitive Decline in Older Persons. Sleep 36, 1027–1032 (2013).

3. Musiek, E. S. & Holtzman, D. M. Mechanisms linking circadian clocks, sleep, and neurodegeneration. Science (80-.). 354, 1004–1008 (2016).

4. Baril, A. A. et al. Biomarkers of dementia in obstructive sleep apnea. Sleep Med. Rev. 42, 139–148 (2018).

5. Elias, A. et al. Risk of Alzheimer’s disease in obstructive sleep apnea syndrome: Amyloid-β and tau imaging. J. Alzheimer’s Dis. 66, 733–741 (2018).

6. Mander, B. A. et al. β-amyloid disrupts human NREM slow waves and related hippocampus-dependent memory consolidation. Nat. Neurosci. 18, 1051–1057 (2015).

7. Ju, Y. S. et al. Slow wave sleep disruption increases cerebrospinal fluid amyloid-b levels. Brain 140, 2104–2111 (2017).

8. Lucey, B. P. et al. Reduced non–rapid eye movement sleep is associated with tau pathology in early Alzheimer’s disease. Sci. Transl. Med. 11, eaau6550 (2019).

9. Kang, J.-E. et al. Amyloid-ß dynamics are regulated by orexin and the sleep-wake cycle. Science (80-.). 326, 1005–1007 (2009).

10. Ju, Y. E. S. et al. Sleep quality and preclinical Alzheimer disease. JAMA Neurol. 70, 587–593 (2013).

11. Sprecher, K. E. et al. Poor sleep is associated with CSF biomarkers of amyloid pathology in cognitively normal adults. Neurology 89, 445–453 (2017).

12. Lucey, B. P. et al. Effect of sleep on overnight cerebrospinal fluid amyloid β kinetics. Ann. Neurol. 83, 197–204 (2018).

13. Ooms, S. et al. Effect of 1 night of total sleep deprivation on cerebrospinal fluid β-amyloid 42 in healthy middle-aged men a randomized clinical trial. JAMA Neurol. 71, 971–977 (2014).

14. Van Egroo, M. et al. Sleep–wake regulation and the hallmarks of the pathogenesis of Alzheimer’s disease. Sleep 42, 1–13 (2019).

15. Ju, Y. E. S., Lucey, B. P. & Holtzman, D. M. Sleep and Alzheimer disease pathology-a bidirectional relationship. Nat. Rev. Neurol. 10, 115–119 (2014).

16. Wang, C. & Holtzman, D. M. Bidirectional relationship between sleep and Alzheimer’s disease: role of amyloid, tau, and other factors. Neuropsychopharmacology (2019). doi:10.1038/s41386-019-0478-5

17. Ulrich, D. Sleep Spindles as Facilitators of Memory Formation and Learning. Neural Plast. 2016, (2016).

18. Gais, S., Mölle, M., Helms, K. & Born, J. Learning-dependent increases in sleep spindle density. J. Neurosci. 22, 6830–6834 (2002).

19. Schmidt, C. et al. Encoding difficulty promotes postlearning changes in sleep spindle activity during napping. J. Neurosci. 26, 8976–8982 (2006).

20. Mednick, S. et al. The critical role of sleep spindles in hippocampal-dependent memory: a pharmacology study. J. Neurosci. 33, 4494–4504 (2013).

21. Miyamoto, D., Hirai, D. & Murayama, M. The roles of cortical slow waves in synaptic plasticity and memory consolidation. Front. Neural Circuits 11, 1–8 (2017).

22. Helfrich, R. F., Mander, B. A., Jagust, W. J., Knight, R. T. & Walker, M. P. Old Brains Come Uncoupled in Sleep: Slow Wave-Spindle Synchrony, Brain Atrophy, and Forgetting. Neuron 97, 221–230.e4 (2018).

23. Winer, J. R. et al. Sleep as a potential biomarker of tau and β-amyloid burden in the human brain Abbreviated Title Authors Center for Human Sleep Science, Department of Psychology, University of California Berkeley, Department of Psychiatry and Human Behavior, Universit. (2019).

24. Winer, J. R. et al. Sleep Disturbance Forecasts β-Amyloid Accumulation across Subsequent Years. Curr. Biol. 30, 4291–4298.e3 (2020).

25. Achermann, P. & Borbély, A. A. Low-frequency (< 1 hz) oscillations in the human sleep electroencephalogram. Neuroscience 81, 213–222 (1997).

26. Lee, J., Kim, D. & Shin, H. S. Lack of delta waves and sleep disturbances during non-rapid eye movement sleep in mice lacking α1G-subunit of T-type calcium channels. Proc. Natl. Acad. Sci. U. S. A. 101, 18195–18199 (2004).

27. Hubbard, J. et al. Rapid fast-delta decay following prolonged wakefulness marks a phase of wake-inertia in NREM sleep. Nat. Commun. 11, 1–16 (2020).

28. Bouchard, M. et al. Sleeping at the switch. Elife 10, (2021).

29. Dang-Vu, T. T. et al. Cerebral correlates of delta waves during non-REM sleep revisited. Neuroimage 28, 14–21 (2005).

30. Dang-Vu, T. T. et al. Functional neuroimaging insights into the physiology of human sleep. Sleep 33, 1589–1603 (2010).

31. Saletin, J. M., van der Helm, E. & Walker, M. P. Structural brain correlates of human sleep oscillations. Neuroimage 83, 658–668 (2013).

32. Stark, S. M., Yassa, M. A., Lacy, J. W. & Stark, C. E. L. A task to assess behavioral pattern separation (BPS) in humans: Data from healthy aging and mild cognitive impairment. Neuropsychologia 51, 2442–2449 (2013).

33. Marks, S. M., Lockhart, S. N., Baker, S. L. & Jagust, W. J. Tau and β-amyloid are associated with medial temporal lobe structure, function, and memory encoding in normal aging. J. Neurosci. 37, 3192–3201 (2017).

34. Grothe, M. J. et al. In vivo staging of regional amyloid deposition. Neurology 89, 2031–2038 (2017).

35. Achermann, P., Dijk, D. J., Brunner, D. P. & Borbély, A. A. A model of human sleep homeostasis based on EEG slow-wave activity: Quantitative comparison of data and simulations. Brain Res. Bull. 31, 97–113 (1993).

36. Fernandez, L. M. J. & Lüthi, A. Sleep spindles: Mechanisms and functions. Physiol. Rev. 100, 805–868 (2020).

37. Adamantidis, A. R., Gutierrez Herrera, C. & Gent, T. C. Oscillating circuitries in the sleeping brain. Nat. Rev. Neurosci. 20, 746–762 (2019).

38. Ju, Y.-E. S., Lucey, B. P. & Holtzman, D. M. Sleep and Alzheimer disease pathology - a bidirectional relationship. Nat Rev Neurol 10, 115–9 (2015).

39. Xie, L. et al. Sleep drives metabolite clearance from the adult brain. Science (80-.). 342, 373–377 (2013).

40. Van Egroo, M. et al. Preserved wake-dependent cortical excitability dynamics predict cognitive fitness beyond age-related brain alterations. Commun. Biol. 2, 1–10 (2019).

41. Chylinski, D. et al. Heterogeneity in the links between sleep arousals, amyloid-b, and cognition. JCI Insight 6, e152858 (2021).

42. Mölle, M., Bergmann, T. O., Marshall, L. & Born, J. Fast and slow spindles during the sleep slow oscillation: Disparate coalescence and engagement in memory processing. Sleep 34, 1411–1421 (2011).

43. Rizzolo, L. et al. Relationship between brain AD biomarkers and episodic memory performance in healthy aging. Brain Cogn. 148, (2021).

44. Berthomier, C. et al. Automatic Analysis of Single-Channel Sleep EEG : Validation in Healthy Individuals. Sleep 30, 1587–1595 (2007).

45. Peter-Derex, L. et al. Automatic analysis of single-channel sleep EEG in a large spectrum of sleep disorders. J. Clin. Sleep Med. (2020). doi:https://doi.org/10.5664/jcsm.8864

46. Chylinski, D. et al. Validation of an Automatic Arousal Detection Algorithm for Whole-Night Sleep EEG Recordings. Clocks & Sleep 2, 258–272 (2020).

47. Coppieters ‘t Wallant, D. et al. Automatic artifacts and arousals detection in whole-night sleep EEG recordings. J. Neurosci. Methods 258, 124–133 (2016).

48. Rosinvil, T. et al. Are age and sex effects on sleep slow waves only a matter of EEG amplitude? Sleep 1–33 (2020). doi:10.1093/sleep/zsaa186

49. Gaudreault, P. O. et al. The association between white matter and sleep spindles differs in young and older individuals. Sleep 41, 1–13 (2018).

50. Lafortune, M. et al. Sleep spindles and rapid eye movement sleep as predictors of next morning cognitive performance in healthy middle-aged and older participants. J. Sleep Res. 23, 159–167 (2014).

51. Martin, N. et al. Topography of age-related changes in sleep spindles. Neurobiol. Aging 34, 468–476 (2013).

52. Weiskopf, N. & Helms, G. Multi-parameter mapping of the human brain at 1mm resolution in less than 20 minutes. Proc. Int. Soc. Magn. Reson. Med. 16, 2241 (2008).

53. Tabelow, K. et al. hMRI – A toolbox for quantitative MRI in neuroscience and clinical research. Neuroimage 194, 191–210 (2019).

54. Ashburner, J. & Friston, K. J. Unified segmentation. Neuroimage 26, 839–851 (2005).

55. Ashburner, J. A fast diffeomorphic image registration algorithm. Neuroimage 38, 95–113 (2007).

56. Tzourio-Mazoyer, N. et al. Automated anatomical labeling of activations in SPM using a macroscopic anatomical parcellation of the MNI MRI single-subject brain. Neuroimage 15, 273–289 (2002).

57. Klunk, W. E. et al. The Centiloid project: Standardizing quantitative amyloid plaque estimation by PET. Alzheimer’s Dement. 11, 1–15.e4 (2015).

58. Narbutas, J. et al. Associations between Cognitive Complaints, Memory Performance, Mood, and Amyloid-β Accumulation in Healthy Amyloid Negative Late-Midlife Individuals. J. Alzheimer’s Dis. 83, 127–141 (2021).

59. Stark, S. M., Stevenson, R., Wu, C., Rutledge, S. & Stark, C. E. L. Stability of age-related deficits in the mnemonic similarity task across task variations. Behav. Neurosci. 129, 257–268 (2015).

60. Jaeger, B. C., Edwards, L. J., Das, K. & Sen, P. K. An R2statistic for fixed effects in the generalized linear mixed model. J. Appl. Stat. 44, 1086–1105 (2017).

61. Kain, M. P., Bolker, B. M. & McCoy, M. W. A practical guide and power analysis for GLMMs: Detecting among treatment variation in randomeffects. PeerJ 2015, (2015).

62. Faul, F., Erdfelder, E., Buchner, A. & Lang, A. G. Statistical power analyses using G*Power 3.1: Tests for correlation and regression analyses. Behav. Res. Methods 41, 1149–1160 (2009).

